# Drp1 SUMO/deSUMOylation by Senp5 isoforms influences ER tubulation and mitochondrial dynamics to regulate brain development

**DOI:** 10.1101/2021.04.15.439911

**Authors:** Seiya Yamada, Ayaka Sato, Hiroki Akiyama, Shin-ichi Sakakibara

**Affiliations:** Laboratory for Molecular Neurobiology, Faculty of Human Sciences, Waseda University, Tokorozawa, Saitama 359-1192, Japan; Advanced Research Center for Human Sciences, Waseda University

**Keywords:** SUMO, Senp5, Drp1, ER, ubiquitin, neuronal polarization, cortical development

## Abstract

Brain development is a highly orchestrated process requiring spatiotemporally regulated mitochondrial dynamics. Drp1, a key molecule in the mitochondrial fission machinery, undergoes various post-translational modifications including conjugation to the small ubiquitin-like modifier (SUMO). However, the functional significance of SUMOylation/deSUMOylation on Drp1 remains controversial. SUMO-specific protease 5 (Senp5L) catalyzes the deSUMOylation of Drp1. We revealed that a splicing variant of Senp5L, Senp5S, which lacks peptidase activity, prevents deSUMOylation of Drp1 by competing against other Senps. The altered SUMOylation level of Drp1 induced by Senp5L/5S affects Drp1 ubiquitination and tubulation of the endoplasmic reticulum (ER), thereby influencing mitochondrial morphology. A dynamic SUMOylation/deSUMOylation balance controls neuronal polarization and migration during the development of the cerebral cortex. These findings suggest a novel role of post translational modification, in which a deSUMOylation enzyme isoform competitively regulates mitochondrial dynamics and ER tubulation via Drp1 SUMOylation levels in a tightly controlled process of neuronal differentiation and corticogenesis.

## INTRODUCTION

The small ubiquitin-like modifier (SUMO) is a post-translational modifier that influences multiple cellular processes such as protein trafficking, protein stability, transcriptional activation/repression, DNA repair, cell-cycle progression, immune response, and dendritic spinogenesis^1–4^. SUMO are evolutionarily conserved small polypeptides (□10 kDa) in eukaryotes. They are covalently attached to the ε-amino groups of lysine residues in all target proteins^5^. Among the five SUMO paralogs (SUMO1–5) in mammals, SUMO2 and SUMO3 share a 96% identical sequence including a conserved sequence for SUMOylation. Thus, SUMO2 and SUMO3 can form a polySUMO chain on the target protein^6, 7^. SUMOylation is achieved through a cascade of three enzymatic reactions comprising a SUMO-activating enzyme (E1), a conjugating enzyme (E2), and SUMO ligases (E3)^4^. Deconjugation or deSUMOylation is mediated by sentrin/SUMO-specific proteases (Senps).

SUMOylation/deSUMOylation is indispensable for neural development^8–10^. The components of the SUMOylation machinery, including SUMO1–3 and a SUMO-conjugating enzyme E2 (Ubc9), are abundantly expressed in neural stem/progenitor cells (NSPCs) and immature migrating neurons during mouse brain development^11^.

Cell division of NSPCs and subsequent migration of neuronal progenies are strictly controlled spatiotemporally during mammalian brain development. Differentiating neurons are classified into stages 1 to 5 based on their *in vitro* morphological characteristics^12, 13^. Neurons extend filopodia (stage 1) to form multiple immature neurites (stage 2). One of the immature neurites becomes an axon (stage 3), while the others develop into dendrites (stage 4). Dendritic spines are then established (stage 5). In the embryonic cerebral cortex, the newborn neurons migrate radially toward the cortical plate (CP) while undergoing a sequential morphological transformation from multipolar (stages 1 and 2) to bipolar. This occurs within the subventricular zone (SVZ) and lower area of the intermediate zone^14^. The trailing process of a bipolar cell becomes an axon; the multipolar to bipolar transition is a polarity establishment process corresponding to stage 3. The leading processes develop into dendrites after the cell reaches its destination (stage 4). Subsequently, dendritic spines are established (stage 5) to enable the synaptic transmission within neural circuits^14^. Dysregulation or failure of polarity formation in migrating neurons causes severe brain malformation and psychiatric disorders such as epilepsy and mental retardation^14, 15^.

A mitochondrion is a highly dynamic organelle that undergoes fission and fusion. Dynamin-related protein 1 (Drp1) has a crucial role in the mitochondrial fission machinery. Cytosolic Drp1 is recruited to the outer mitochondrial membrane and oligomerizes to activate GTP-dependent mitochondrial fission^16, 17^. A Drp1 deficiency impedes mitochondrial fission and induces mitochondrial aggregates, which disturb proper neural development, neuronal fate commitment, neurite extension, dendrite development, and synapse formation^18, 19^. In humans, the dominant-negative allele of the *DRP1* gene causes a broad range of abnormalities such as brain malformation and optic atrophy, leading to early neonatal death^20^. These findings establish the importance of Drp1-mediated regulation of mitochondrial dynamics in brain development.

SUMOylation/deSUMOylation controls mitochondrial dynamics through Drp1 modulation. SUMOylation contributes to Drp1 stability, whereas deSUMOylation decreases it^21–23^. Further, Drp1 localizes to the lumen of the endoplasmic reticulum (ER)^17^, and ER-associated Drp1 induces ER tubulation. ER tubules mechanically contact the mitochondria, which facilitates the recruitment of cytosolic Drp1 to the contact sites. This results in mitochondrial division^17^.

The mechanism of Drp1-mediated mitochondrial fission is established^24^; however, the mechanism of Drp1 SUMOylation-mediated mitochondrial dynamics remains unclear ^8, 25^. Moreover, we find no reports showing the detailed machinery for Sumoylation-dependent regulation of brain development. Here, we report that the novel Senp5 isoform, which lacks peptidase activity, prevents deSUMOylation in a competitive manner. We further demonstrated *in vitro* and *in utero* its involvement in neuronal development during corticogenesis.

## RESULTS

### Identification of the Senp5S isoform

Previous studies reported that the mouse *Senp5* gene encoded a single 749 amino acid (aa)-protein with a molecular mass of 97 kDa (PMID: 25128087) (Fig. 1a)^26^. However, our database search of public cDNA sequences (GenBank, and dbEST containing the single pass cDNA sequences and Expressed Sequence Tags) predicted the existence of other *Senp5* isoforms that lack the C-terminal catalytic domain. We performed RT-PCR using mRNA isolated from E12 and adult mouse brains and confirmed the expression of *Senp5 short* isoforms, which were generated by alternative splicing of *Senp5* transcript. We designated one of these short isoforms as Senp5S and the long isoform as Senp5L (Fig. 1a). Senp5S was predicted to have a molecular mass of 72 kDa and lack the enzymatic catalytic center (Cys 707) critical for peptidase activity. A comparable short isoform of Senp5 lacking most of the residues of the C-terminal catalytic triad is known in humans. Quantitative PCR (qPCR) analysis indicated that *Senp5L* and *Senp5S* mRNAs are concurrently expressed during embryonic stages and gradually upregulated throughout brain development (Fig. 1b, c). Consistently, western blot analysis with anti-Senp5 antibody revealed the developmentally regulated expression of Senp5 isoform proteins (Fig. 1d). Senp5L protein with a size of ∼100 kDa was detected throughout brain development. However, Senp5S was barely detectable in the early embryonic brain (E12) but rapidly increased in the late embryonic and postnatal periods as doublet bands of ∼75 kDa (Fig. 1d). Western blotting of cultured NSPCs and differentiating cortical neurons cultured for 3 days *in vitro* (div) showed that NSPCs solely express Senp5L, whereas immature neurons start to express Senp5S (Fig. 1d). We concluded that Senp5L is a dominant form in the early embryonic stages E12□E14 when active neurogenesis occurs, while Senp5S becomes upregulated during the differentiation stage and becomes the dominant form in postnatal and adult brains.

**Figure 1.**
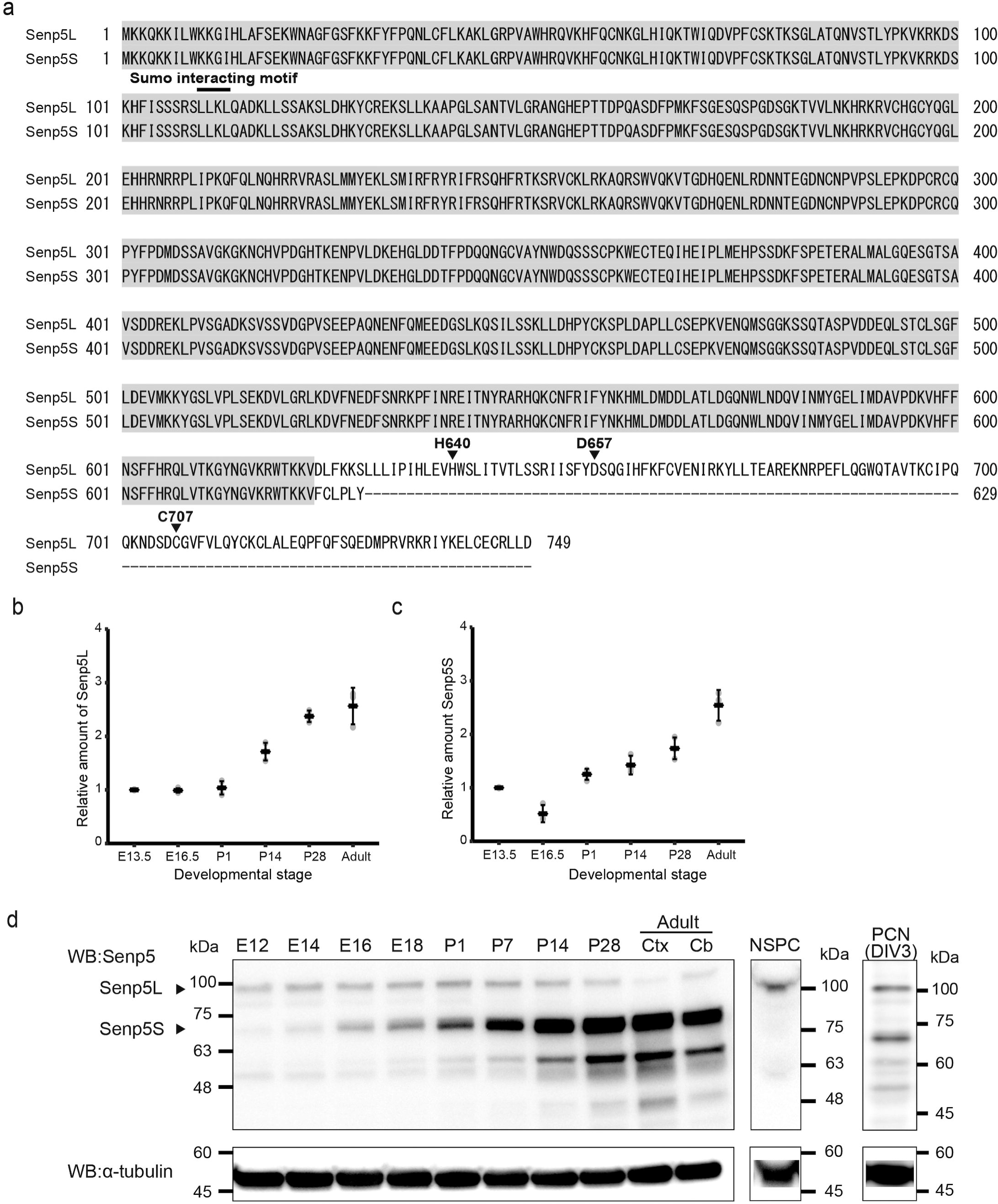
Identification of Senp5S isoform. (a) Shown are the deduced amino acid sequences of mouse Senp5L and Senp5S. Identical amino acids are highlighted. Gaps in the alignment are indicated by dashes. Arrowhead denotes the catalytic center Cys-707 necessary for peptidase activity. His-640, Asp-657, and Cys-707 constitute the catalytic triad for cysteine protease. The N-terminal LLKL sequence (110–113) is the SUMO-interacting motif. Only Senp5S lacks the C-terminal catalytic center, though both isoforms have the N-terminal SUMO-interacting motif. (b, c) Developmental changes in *Senp5L* (b) and *Senp5S* (c) mRNA expression in the mouse cerebral cortex by qPCR analysis. Gray dots represent three independent experiments normalized to the corresponding β-actin mRNA. Mean ± SD are also shown. (d) Developmental changes in Senp5L/5S protein expression in the mouse brain. Total protein extracts were prepared from the whole brains (E12–P28), adult cerebral cortex (ctx), or adult cerebellum (cb), and subjected to immunoblotting with anti-Senp5 antibody (left panel). Protein lysate from primary cultured neural stem/precursor cells (NSPC) or primary cultured cortical neurons (PCN) at 3 div was also analyzed by immunoblot (middle and right panels). The blots were reprobed with anti-α-tubulin antibody (bottom panels) to examine protein loading quantitatively.

### Senp5S promotes global SUMOylation

To test whether Senp5S affects protein deSUMOylation, we monitored global SUMOylation in HEK293T cells expressing enhanced green fluorescent protein fused to Senp3 (EGFP-Senp3), -Senp5L, or -Senp5S and hemagglutinin (HA)-SUMO3 or HA-SUMO1. In control EGFP-expressing cells, many HA-SUMO3-conjugated proteins were detected as ladder bands with high molecular weight (Fig. 2a). These SUMO3-conjugated proteins were dramatically reduced or even diminished by exogenous expression of Senp3 or Senp5L, thus confirming the deSUMOylation activity of Senp3 and Senp5. In contrast, Senp5S substantially increased SUMO3-conjugated proteins compared to the control. This Senp5S-mediated SUMOylation was almost canceled by co-transfection of Senp3 or Senp5L, which suggested the competitive nature of Senp5S and Senp5L. Same results were also observed in Neuro2a cells (data not shown). However, we did not detect a massive accumulation of SUMOylated proteins incorporated with HA-SUMO1 (Fig. 2a). These results indicated that Senp5S inhibited deSUMOylation by Senps, which resulted in the massive accumulation of SUMOylated substrates in cells. Senp5S was likely a competitor against Senp5L.

**Figure 2.**
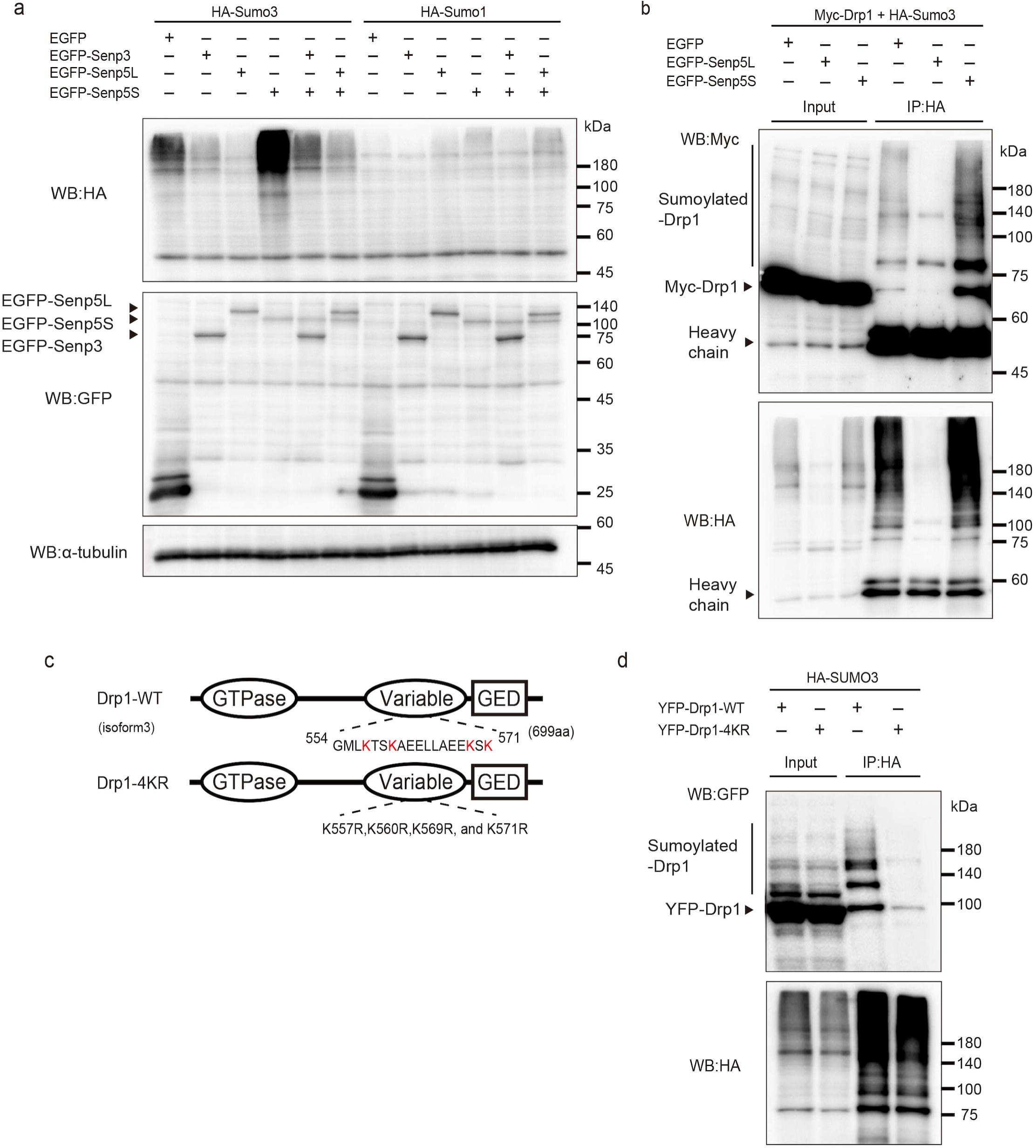
Senp5S competes with other Senps and promotes SUMOylation. (a) Global SUMO3-ylation levels are changed by expression of Senp5 isoforms. HEK293T cells were transfected with EGFP, EGFP-Senp3, EGFP-Senp5L, or EGFP-Senp5S along with HA-SUMO3 or HA-SUMO1, followed by immunoblotting with anti-HA (upper panel), anti-GFP (middle panel), or anti-α-tubulin (bottom panel). (b) Effect of Senp5L and Senp5S expression on SUMO3-ylation of Drp1. HEK293T expressing EGFP, EGFP-Senp5L, or EGFP-Senp5S along with Myc-Drp1 and HA-SUMO3 were immunoprecipitated (IP) with anti-HA antibody. Total cell lysates (input) or IP samples were analyzed by immunoblotting with anti-Myc (upper panel) and anti-HA (bottom panel). SUMOylated-Drp1 is indicated by the perpendicular line. Heavy chain, immunoglobulin heavy chain of the HA antibody. (c) Domain structures of wild-type Drp1 (Drp1-WT) and a non-SUMOylatable Drp1 mutant (Drp1-4KR). The D-octadecapeptide sequence, comprising 18 amino acid residues (554-571) encompassing in the variable domain, is indicated. Drp1-4KR was created by substituting four SUMO-acceptor lysine residues (*red*) in the D-octadecapeptide with arginine residues. Oval and rectangle indicate GTPase activity domain and GTPase effector domain (GED), respectively. (d) Drp1 was SUMO3-ylated through the D-octadecapeptide sequence. HEK293 cells were co-transfected with HA-SUMO3, p14-Arf, Ubc9, and YFP-Drp1-WT or YFP-Drp1-4KR, then immunoprecipitated with anti-HA. Cell lysates (input) or IP samples were analyzed by immunoblotting with anti-GFP (top panel) or anti-HA (bottom panel). SUMOylated-Drp1 is indicated by the perpendicular line.

### Senp5 isoforms control Drp1 deSUMO/SUMOylation

There has been no evidence of the functional significance of Drp1 SUMOylation with SUMO3 in mitochondrial dynamics. However, it is known that Drp1 SUMOylation by SUMO1 enhances Drp1 stabilization and directs mitochondrial fragmentation^21, 22, 25^. We investigated Drp1 SUMO3-conjugation and the role of Senp5 isoforms. Co-immunoprecipitation (IP) showed SUMO3-ylation of myc-Drp1 and Senp5L-mediated deconjugation, as previously described (Fig. 2b)^22^. Global SUMO3-ylation was greatly reduced in the cell lysate expressing EGFP-Senp5L. Interestingly, Senp5S clearly enhanced SUMO3-ylation of Drp1 compared to the control (Fig. 2b). SUMO3-ylation of Drp1 was confirmed by co-IP using non-SUMOylatable Drp1, in which four acceptor lysine residues encompassing the variable domain were substituted with arginine (Drp1-4KR)^22, 25^ (Fig. 2c). As shown in Fig. 2d, we observed no SUMO3-conjugation with Drp1-4KR. These results showed that Drp1 is a substrate for SUMO3-ylation. Furthermore, Senp5L and Senp5S have opposite effects on Drp1 SUMOylation.

### Senp5 isoforms alter mitochondrial dynamics

A previous study reported that overexpression of Senp5 in Hela cells induced mitochondrial elongation^23^. Thus, we examined whether Senp5S affected mitochondrial morphology. To visualize the shape of individual mitochondria, we introduced into HEK293T cells the reporter plasmid mitochondria-targeted monomeric Kusabira-Orange1 (pMT-mKO1), Senp5L, and Senp5S. Consistent with the previous study, Senp5L expression increased mitochondrial length compared to the control (Fig. 3a). In contrast, Senp5S caused mitochondrial fragmentation, resulting in shortened mitochondria. SUMOylation-dependent fragmentation of mitochondria was further supported by the effect of overexpression of SUMO3 or SUMO1 during mitochondrial fragmentation (Fig. 3b, Supplementary Fig. 1). SUMO3-induced fragmentation was further increased by Senp5S co-expression and decreased by Senp5L co-expression (Fig. 3b). These observations strongly indicate that mitochondrial dynamics are regulated, at least partly, by the SUMO3-ylation level of Drp1 in a Senp5L/S dependent manner. To further confirm the involvement of Drp1-SUMOylation in mitochondrial fission, we evaluated the mitochondrial morphology of cells overexpressing Drp1-wild type (WT) or Drp1-4KR. As expected, mitochondrial fragmentation was accelerated by Drp1-WT but not by Drp1-4KR (Fig. 4d, e). We conclude that Senp5L and Senp5S promote mitochondrial fusion and fission, respectively, via the regulation of Drp1 deSUMO/SUMOylation. Considering the competitive nature of Senp5 isoforms, the expression levels of Senp5L and Senp5S might be critical for mitochondrial dynamics.

**Figure 3.**
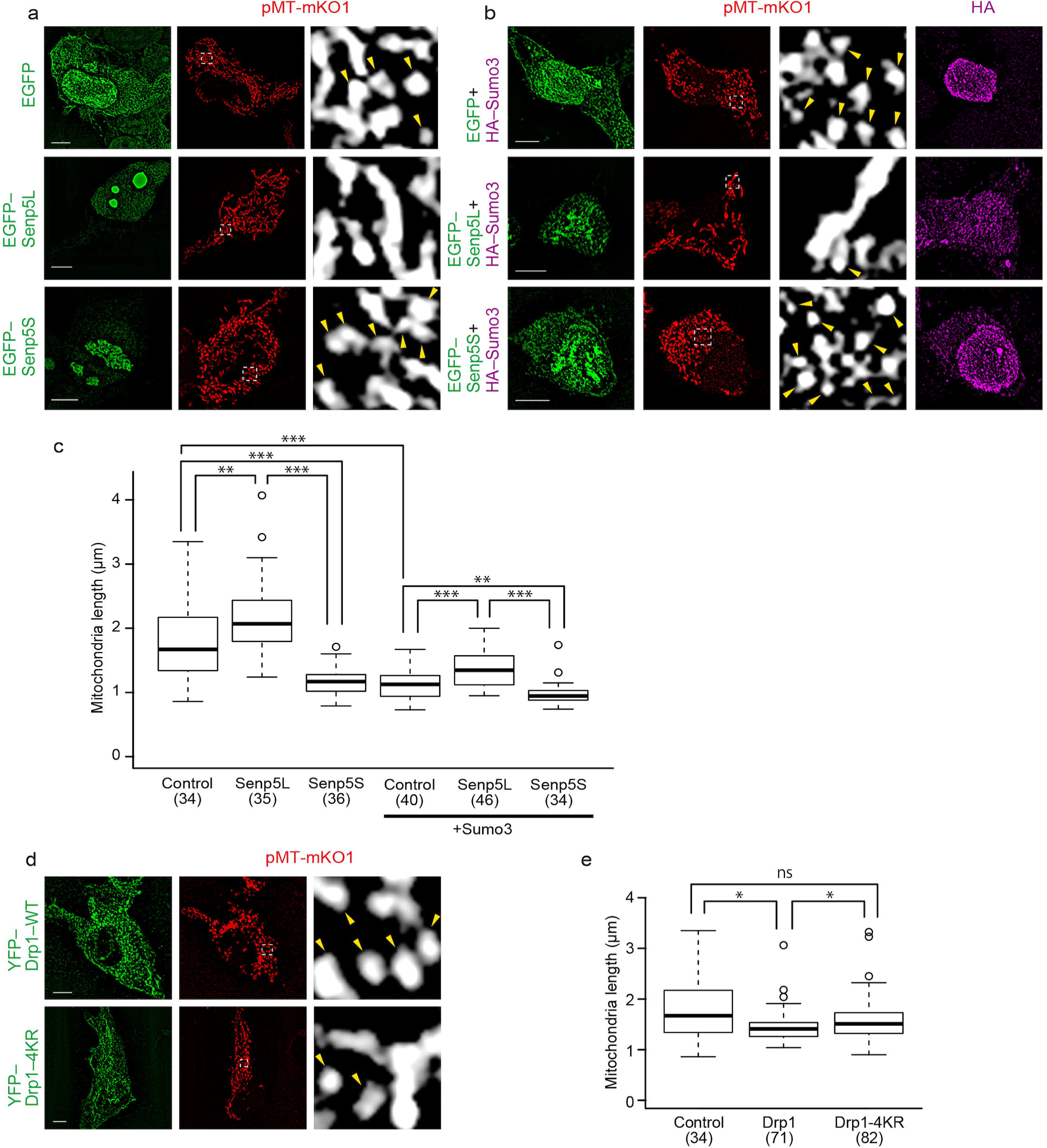
Senp5L/5S and Drp1-SUMOylation regulate mitochondrial morphology. (a) Mitochondrial morphologies in 293T cells expressing EGFP, EGFP-Senp5L, or EGFP-Senp5S were analyzed after deconvolution of the confocal projection images. pMT-mKO1 (*red*) was co-transfected to visualize mitochondria. Right column is higher magnification of the boxed areas, depicting individual mitochondrial morphology (*white*). Arrowheads denote fragmented mitochondria. (b) Overexpression of SUMO3 enhances the mitochondrial fragmentation. Cells were co-transfected with HA-SUMO3 along with EGFP, EGFP-Senp5L, or EGFP-Senp5S. HA-SUMO3 expression was confirmed by immunostaining with anti-HA (*magenta*). (c) Quantified comparison of the effect of Senp5L, Senp5S, and SUMO3 overexpression on mitochondrial length. Box and whisker plots summarize the length (μm) of mitochondria. Numbers in parentheses indicate the numbers of cells examined in multiple independent experiments in (a) and (b). **, P < 0.01; ***, P < 0.001; Welch’s t-tests with Holm-Bonferroni correction. (d) Confocal projection images of HEK293T cells transfected with YFP-Drp1-WT or YFP-Drp1-4KR (*green*). pMT-mKO1 (*red*) was co-transfected to visualize mitochondria. Right column: magnified views of the boxed areas showing individual mitochondria (*white*). Arrowheads indicate fragmented mitochondria. (e) Quantification of mitochondria length in cells expressing Drp1-WT and Drp1-4KR. Box and whisker plots summarize the mitochondrial length (μm). Numbers in parentheses indicate the numbers of cells measured for mitochondrial length determination. ns, not significant, *, P < 0.05; Welch’s t-tests with Holm-Bonferroni correction. Scale bars, 5 μm.

**Figure 4.**
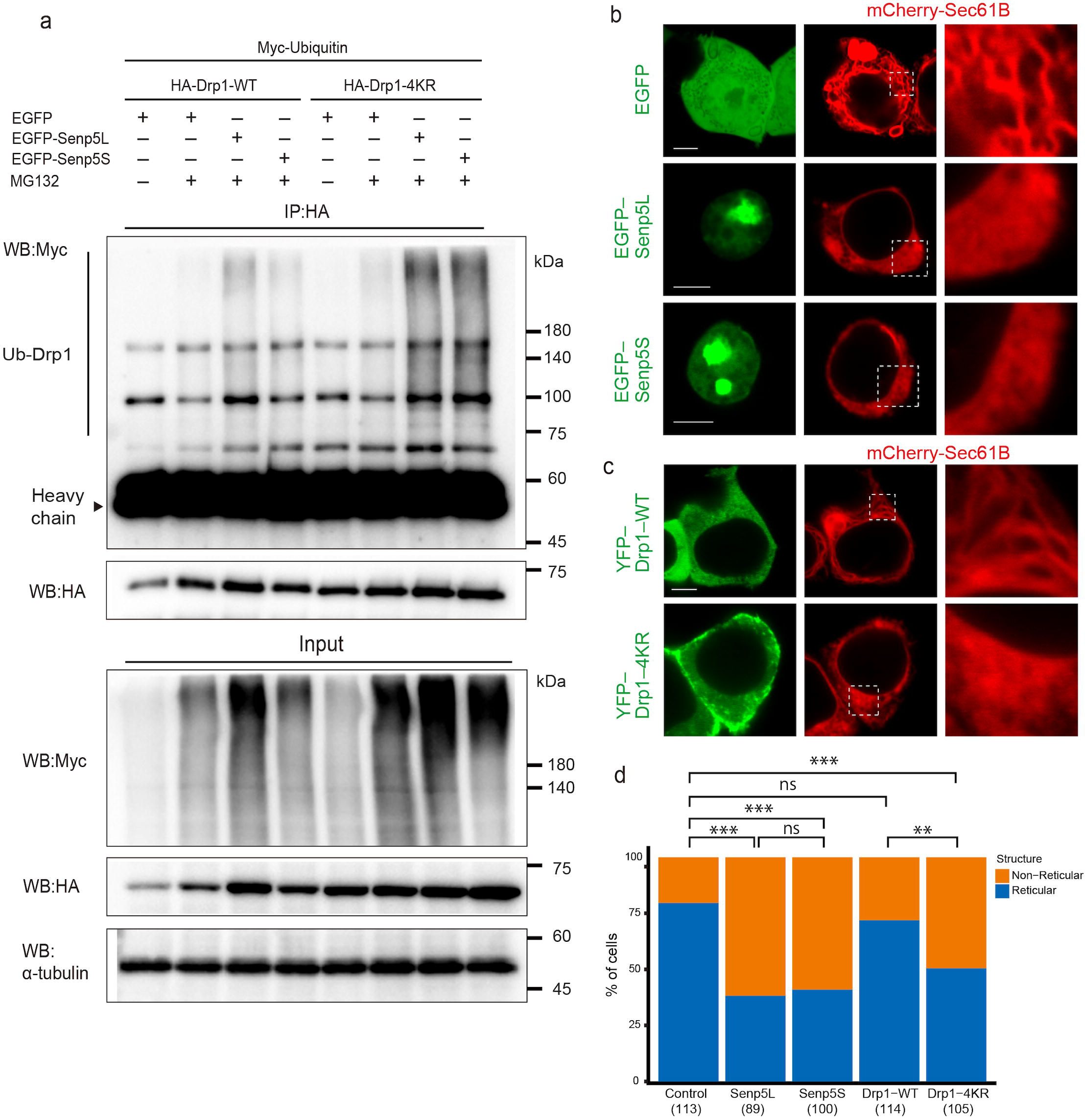
Senp5L/5S are involved in Drp1 ubiquitination and ER dynamics. (a) SUMOylation prevents the ubiquitination of Drp1. HEK293T cells were transfected with Myc-Ubiquitin and HA-Drp1-WT or HA-Drp1-4KR constructs with or without pretreatment with 5 μM MG132 for 3 h. To evaluate the effect of Senp5 on ubiquitination of Drp1, cells are additionally co-transfected with EGFP-Senp5L, EGFP-Senp5S, or (as a control) EGFP. Subsequently, each cell lysate was immunoprecipitated with anti-HA, and subjected to immunoblotting with anti-Myc, anti-HA, or anti-α-tubulin. Panels show representative immunoblots for the input lysate (right panels) and IP samples (left panels). Drp1 conjugated with Myc-ubiquitin is observed as the ladder or smear bands with higher molecular weight (vertical line). Heavy chain, immunoglobulin heavy chain of the HA antibody. (b) Senp5L and 5S induce morphological change in the ER. 293T cells were transfected with EGFP, EGFP-Senp5L, or EGFP-Senp5S (*green*), together with mCherry-Sec61B (*red*) to monitor the ER network. Right column: higher magnification of the square areas showing mCherry-Sec61B ^+^ ER morphology. (c) SUMOylation at the D-octadecapeptide of Drp1 is required for inducing tubulation of the ER. HEK293T cells were transfected with mCherry-Sec61B (*red*) and YFP-Drp1-WT or YFP-Drp1-4KR (*green*). Right column: higher magnification of the square areas. (d) Quantified comparison of the effect of Senp5L, Senp5S, or Drp1-4KR overexpression on ER morphology. The stacked bar chart shows the percentages of cells that exhibit a tubulated reticular ER or a sheet-like, non-reticular ER. Numbers in parentheses indicate the numbers of cells observed in two independent experiments in (b) and (c). ns, not significant; **, P < 0.01; ***, P < 0.001; chi-square tests with Holm-Bonferroni correction. Scale bars, 5 μm.

### Senp5 isoforms control Drp1 ubiquitination and ER morphology

Previous study showed that overexpression of SUMO1 stabilized Drp1, resulting in mitochondrial fragmentation^21^. Although the molecular mechanism of Drp1 modulation by SUMO3 remained unclear, one plausible explanation was that Drp1 conjugation with SUMO3 prevented ubiquitination, thus inhibiting the proteasome-mediated degradation of Drp1. We examined Drp1 ubiquitination in the presence of exogenous Senp5L or -Senp5S and found an increased level of ubiquitinated Drp1 in cells expressing Senp5L with or without treatment with N-ethyl-maleimide (NEM), a cysteine peptidases inhibitor (Supplementary Fig. 2a). To further test whether Drp1 ubiquitination was dependent on SUMOylation, we analyzed the ubiquitination level of Drp1-WT and Drp1-4KR. Cells were transfected with myc-ubiquitin together with HA-Drp1-WT or -4KR, and EGFP-Senp5L or -Senp5S, then extracts were immunoprecipitated with anti-HA antibody. The ubiquitinated Drp1-WT and -4KR were then determined. Drp1-4KR was more ubiquitinated than Drp1-WT when pretreated with MG132, a proteasome inhibitor (Fig. 4a). Noticeably, Drp1-WT ubiquitination was increased when co-expressed with Senp5L. These results indicated that deSUMOylation catalyzed by Senp5L promoted Drp1 ubiquitination, which could lead to decreased Drp1 activity and thus mitochondrial fusion. These findings were consistent with our results on mitochondrial length (Fig. 3). Further, we found that less ubiquitination of Drp1-WT was observed in Senp5S- compared to Senp5L-expressing cells (Fig. 4a). Another possible explanation for the involvement of SENP5 in mitochondrial dynamics was an influence of Senp5 on Drp1-dependent regulation of ER structure. A recent study reported that four lysine residues in the variable domain of Drp1 play a critical role in ER tubulation and mitochondrial fragmentation^17^. These four lysine residues are identical to the SUMOylation sites (K557, K560, K569, and K571) (Fig. 2c). Thus, we hypothesized that Drp1 SUMO/deSUMOylation regulates ER tubulation. To test this idea, we examined the ER architecture in HEK293T cells expressing Senp5L or Senp5S. ER membranes were visualized using ER-resident mCherry-Sec61B. As shown in Fig. 4b, tubulated and reticular ER were observed in EGFP-control cells, whereas exogenous expression of Senp5L or Senp5S significantly decreased reticular ER. To test the involvement of Drp1 SUMOylation in ER tubulation, Drp1-WT or the Drp1-4KR construct were introduced into HEK293T cells. Drp1-4KR but not Drp1-WT disrupted the reticular ER structure (Fig. 4c, d). These observations support the conjecture that SUMOylated Drp1 molecules elicit ER tubulation. Considering the suppression of ER tubulation by Senp5S, stringent control of deSUMOylation/SUMOylation by Senp5L and Senp5S seems essential for Drp1-mediated ER tubulation.

### Senp5 expression during brain neurogenesis

To explore the *in vivo* function of the Senp5 isoforms, we examined the Senp5 expression profile in embryonic cerebral cortices at different stages of neurogenesis and the ensuing cell differentiation. Senp5 antibody used in this study recognizes both Senp5L and S proteins. At E13.5, Senp5 immunoreactivity was observed throughout the neocortex including the VZ surrounding the lateral ventricles, where many NSPCs undergo cell division to generate immature neurons (Fig. 5a). Later, at E16.5, dense Senp5 immunoreactivity was observed in the CP, while a substantial level of expression continued in the VZ, SVZ, and IZ (Fig. 5b, c). These immunohistochemical data showed persistent Senp5 expression in neuronal lineage. The expression profile of Senp5 was comparable with that of Senp3 in the developing cortex (Supplementary Fig. 4a–c).

**Figure 5.**
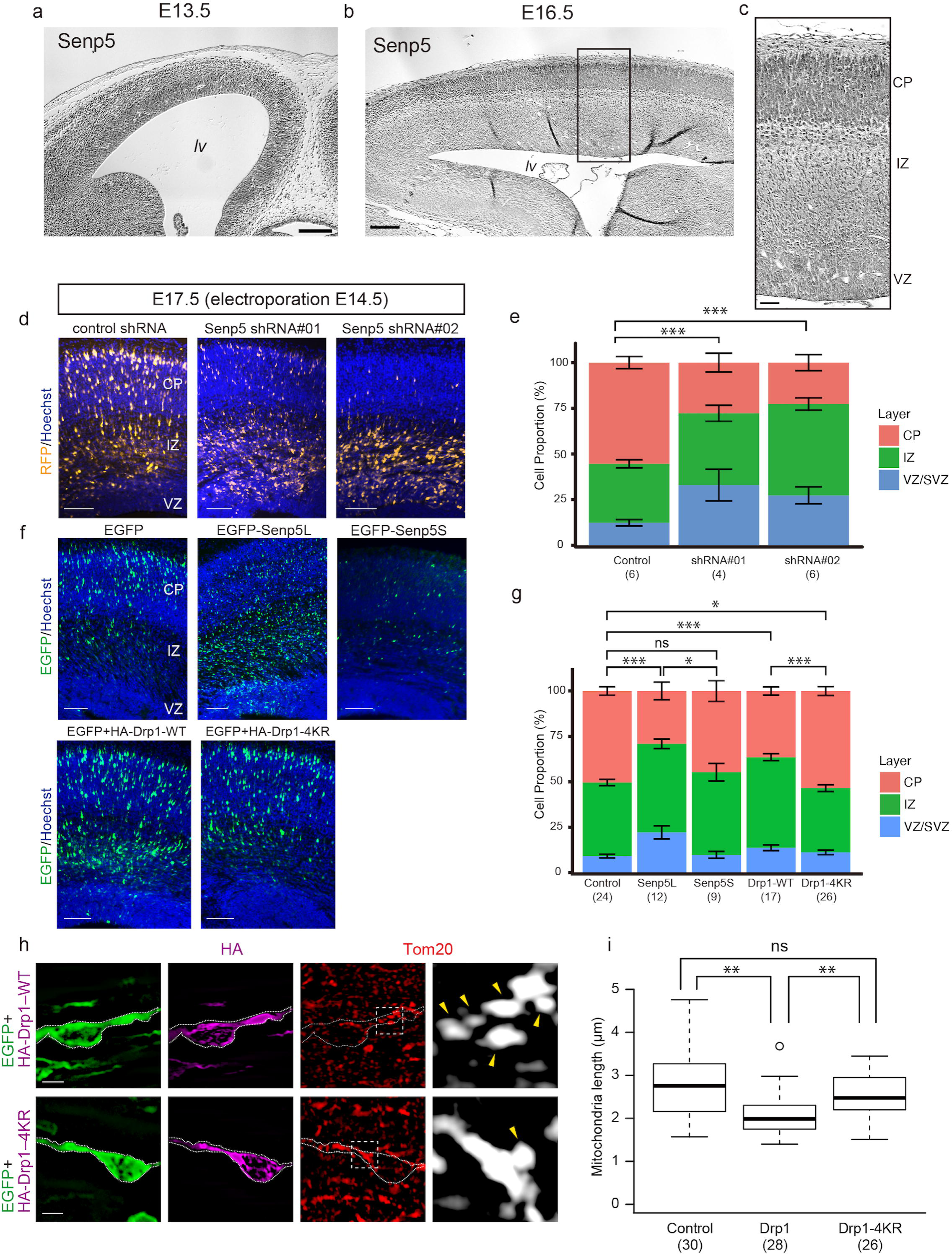
Dysregulation of Senp5L/5S expression or Drp1-SUMOylation cause migration defects of neurons in embryonic cerebral cortex. (a–c) Distribution of Senp5 in embryonic cortex. Coronal sections of cerebral cortex at E13.5 (a) and E16.5 (b) were immunostained with anti-pan-Senp5 antibody. (c) Higher magnification of the boxed area of (b). lv, lateral ventricle; CP, cortical plate; IZ, intermediate zone; VZ, ventricular zone. (d, e) Senp5 knockdown (KD) induces accumulation of migrating neurons in the IZ. Non-targeting control shRNA or Senp5 shRNA construct (shRNA #01 or shRNA #02) was electroporated into E14.5 cortices *in utero* together with TurboRFP. (d) Coronal sections of E17.5 cortex showing the distribution of TurboRFP^+^ cells expressing shRNA (*yellow*). Nuclei were counterstained with Hoechst dye (*blue*). (e) Distribution of TurboRFP^+^ cells in the indicated areas at E17.5. ***, P < 0.001; two-way ANOVA with Holm-Bonferroni correction. (f, g) Overexpression of Senp5L or Drp1-WT represses neuronal migration. (f) EGFP-Senp5L, EGFP-Senp5S, HA-Drp1-WT (with EGFP), HA-Drp1-4KR (with EGFP), or control EGFP (*green*) was electroporated *in utero* at E14.5, and the neocortex was analyzed at E17.5. (g) Distribution of electroporated EGFP^+^ cells in the indicated areas. ns, not significant; *, P < 0.05; ***, P < 0.001; two-way ANOVA with Holm-Bonferroni correction. Numbers in parentheses indicate the numbers of embryos analyzed. (h) Magnified view of immature neurons migrating in the IZ of the brain electroporated with HA-Drp1-WT or HA-Drp1-4RK. Sections were immunostained with anti-Tom20 (*red*) and anti-HA (*magenta*). Right: Higher magnification of the boxed areas showing the morphology of Tom20^+^ mitochondria (*white*). Arrows denote fragmented mitochondria. (i) Box and whisker plots summarize the mitochondrial length data (μm). Forced expression of Drp1-WT altered mitochondrial morphology in migrating neurons. Numbers in parentheses indicate the numbers of cells analyzed. ns, not significant; **, P < 0.01; Welch’s t-tests with Holm-Bonferroni correction. Stacked bar charts represent mean ± SEM. Scale bars: 100 μm in (a) and (b), 25 μm in (c), 100 μm in (d) and (f), and 5 μm in (h).

### Senp5 in cerebral cortex development

To gain direct insight into the role of Senp5 isoforms in brain development, we performed *in vivo* knockdown (KD) and overexpression experiments using *in utero* electroporation in mouse embryos. Plasmids encoding the short hairpin RNA (shRNA) were delivered by electroporation into NSPCs in E14.5 embryos through the lateral ventricle, and brain sections were prepared three days later (E17.5). TurboRFP was electroporated together with shRNA as a reporter. Because of the technical difficulty of designing effective shRNAs specific to Senp5S, we used shRNA constructs (shRNA#01 and #02) that knocked down both Senp5L and 5S (Supplementary Fig. 5a)^27^. In control embryos electroporated with scrambled shRNA, new-born turboRFP^+^ neurons radially migrated away from the VZ toward the pial surface. Numerous turboRFP^+^ cells reached the CP (Fig. 5d). By contrast, Senp5 KD hindered cell migration into the CP. TurboRFP^+^ cells were found in the IZ and VZ/SVZ (% of cells in CP: control, 55.4 ± 3.3%, n = 6; SENP5 shRNA #01, 27.8 ± 5.2%, n = 4; SENP5 shRNA #02, 22.6 ± 4.4%, n = 6; Fig. 5e). These results indicated a critical role for Senp5 in neuronal migration and corticogenesis.

Next, we assessed the overexpression phenotype of each Senp5 isoform. As shown in Figs. 5f and 5g, EGFP-Senp5L significantly inhibited neuronal migration from the VZ/SVZ (% of cells in CP: EGFP, 50.4 ± 2.9%, n = 24; Senp5L; 29.0 ± 4.8%, n = 12). However, the embryos electroporated with EGFP-Senp5S did not exhibit any deranged cortical organization or impaired neuronal migration (% of cells in CP: 44.7 ± 5.7%, n = 9; Fig. 5g). Notably, the number of EGFP-Senp5S^+^ cells in specimens was extremely low (Fig. 5f) compared to that of Senp5L or control EGFP. Considering the phenotypic equivalence between KD and overexpression of Senp5L, we conclude that a strictly controlled level of Senp5L expression must be required for cortical development

### Control of Drp1 SUMOylation level appropriate for cortical development

To examine whether Drp1 SUMOylation level influenced brain development, HA-Drp1-WT or HA-Drp1-4KR^28^ was electroporated *in utero* into E14.5 brains and the brains were analyzed at E17.5. As shown in Fig. 5f, overexpression of Drp1-WT suppressed neuronal migration into CP (% of cells in CP: 36.4 ± 2.3%, IZ: 49.9 ± 1.9%, n = 17). However, Drp1-4KR did not show any apparent defects in cortical organization (CP: 53.5 ± 2.4%, IZ: 35.3 ± 1.9%, n= 26), compared to control (Fig. 5f, g). These results showed that aberrant cortical development caused by Drp1-WT was dependent on the SUMOylation of Drp1. We further investigated whether Drp1 SUMOylation affected mitochondrial dynamics by anti-Tom20 immunostaining of neurons in the IZ that were extending the leading process and acquiring polarization. Consistent with the results obtained with HEK293T cells (Fig. 3d, e), Drp1-WT overexpression shortened the average mitochondrial length of migrating neurons, while Drp1-4KR expression abolished this fragmentation phenotype (Fig. 5h, i). Dysregulated expression of Senp5 induced inappropriate Drp1-SUMOylation. Under such conditions, control of mitochondrial fission/fusion was lost, which resulted in impaired motility of immature neurons.

### Senp5 isoforms regulate neuronal polarization

Neuronal polarization can occur simultaneously with cortical neuronal migration^14^. To elucidate whether Senp5 isoforms are involved in polarity formation in migrating neurons, we performed Senp5 overexpression and KD experiments using a culture of dissociated cortical neurons prepared from E16-17 embryos. TurboRFP was co-transfected to visualize processes extending from individual neurons. Anti-SMI312 immunostaining was performed to detect axon formation. By 3 div, most cells transfected with non-targeting control shRNA had established polarity (stage 3; Fig. 6a). In contrast, more than half of the neurons with reduced Senp5 levels remained in unpolarized stages (stage 1/2; Fig. 6a, b). Furthermore, Senp5 KD cells extended a significantly small number of neurites compared to the control cells (Fig. 6c). To assess the role of the Senp5 isoforms individually, cortical neurons were transfected with mCherry-tagged Senp5L or 5S. Expression of exogenous Senp5L and Senp5S caused defects in the acquisition of neuronal polarity (Fig. 6d, e). Furthermore, the number of neurites was significantly less in Senp5L- and Senp5S-expressing neurons than in control neurons (Fig. 6f). These results demonstrated the essential role of Senp5 isoforms in neuronal polarization. Since all experimental conditions, including KD and Senp5 isoform overexpression, elicited similar defects in neurons, the balanced expression of Senp5L and Senp5S appears required for neurite outgrowth and polarity formation.

**Figure 6.**
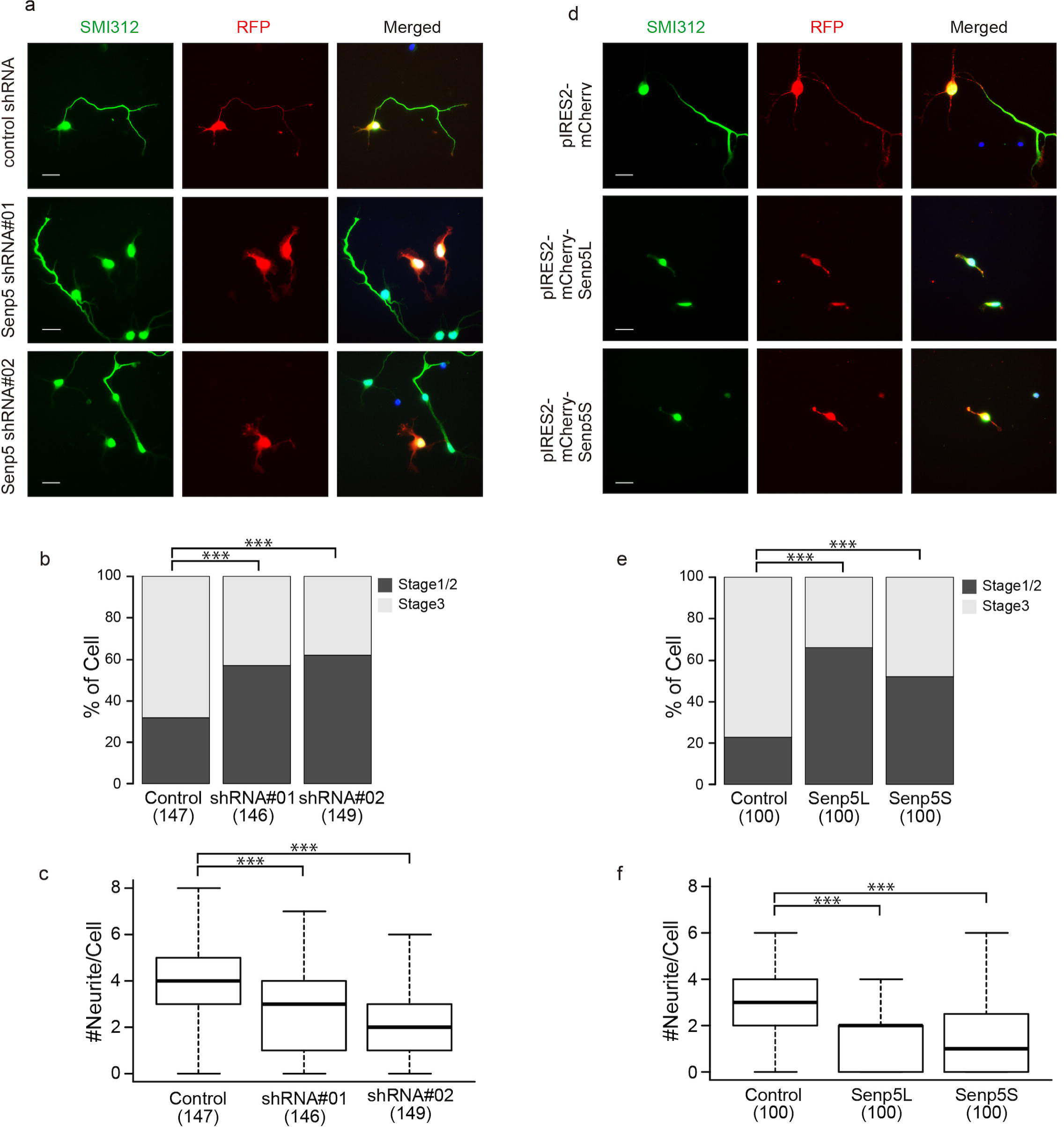
Senp5L/5S regulate neurite outgrowth and polarization. (a–c) Senp5 shRNAs (shRNA #01 or shRNA #02) or control shRNA was introduced into primary cultured cortical neurons along with a reporter plasmid encoding red fluorescent protein (RFP) (*red*). (a) Neurofilaments in individual neurons were stained with anti-SMI312 (*green*) at 3 div. (b) The stacked bar chart shows the percentages of cells having SMI312^+^ axons (stage 3). Numbers in parentheses indicate the numbers of cells analyzed. ***, P < 0.001; chi-square tests with Holm-Bonferroni correction. (c) Box and whisker plots summarize the numbers of RFP^+^ neurites extending from individual neurons. Numbers in parentheses indicate the numbers of cells measured. ns, not significant; ***, P < 0.001; Welch’s *t*-tests with Holm-Bonferroni correction. (d–f) Overexpression of Senp5L and Senp5S in immature neurons. pIRES2-mCherry-Senp5L, pIRES2-mCherry-Senp5S, or control pIRES2-mCherry was electroporated into primary cultured cortical neurons. Neurofilaments and mCherry were stained with anti-SMI312 (*green*) and anti-RFP (*red*) at 3 div. (e) The stacked bar chart shows the percentages of cells having SMI312^+^ axons (stage 3). Numbers in parentheses indicate the numbers of cells measured; ***, P < 0.001; chi-square tests with Holm-Bonferroni correction. (f) Box and whisker plots summarize the numbers of RFP^+^ neurites extending from individual neurons. Numbers in parentheses indicate the numbers of cells measured. ns, not significant; ***, P < 0.001; Welch’s t tests with Holm-Bonferroni correction. Scale bars, 20 μm.

## DISCUSSION

Members of the Senp family (Senp1-8) share a conserved C-terminal catalytic domain that sub serves their cysteine protease activities, while their N-terminal regions have diverse specific functions^29^. All identified Senps have isopeptidase activity, which mediates the deSUMOylation of target proteins. In addition, several Senps have endopeptidase activity, which converts pro-SUMO into conjugatable mature SUMO capable of SUMOylation. Conventional Senps can control both SUMOylation and deSUMOylation depending on protease activity. We revealed a novel protease-independent mechanism for SUMO cycle regulation, in which the newly identified Senp5S, which lacks protease activity, promoted SUMOylation by competing with other Senps for deSUMOylation sites (Fig. 7a).

**Figure 7.**
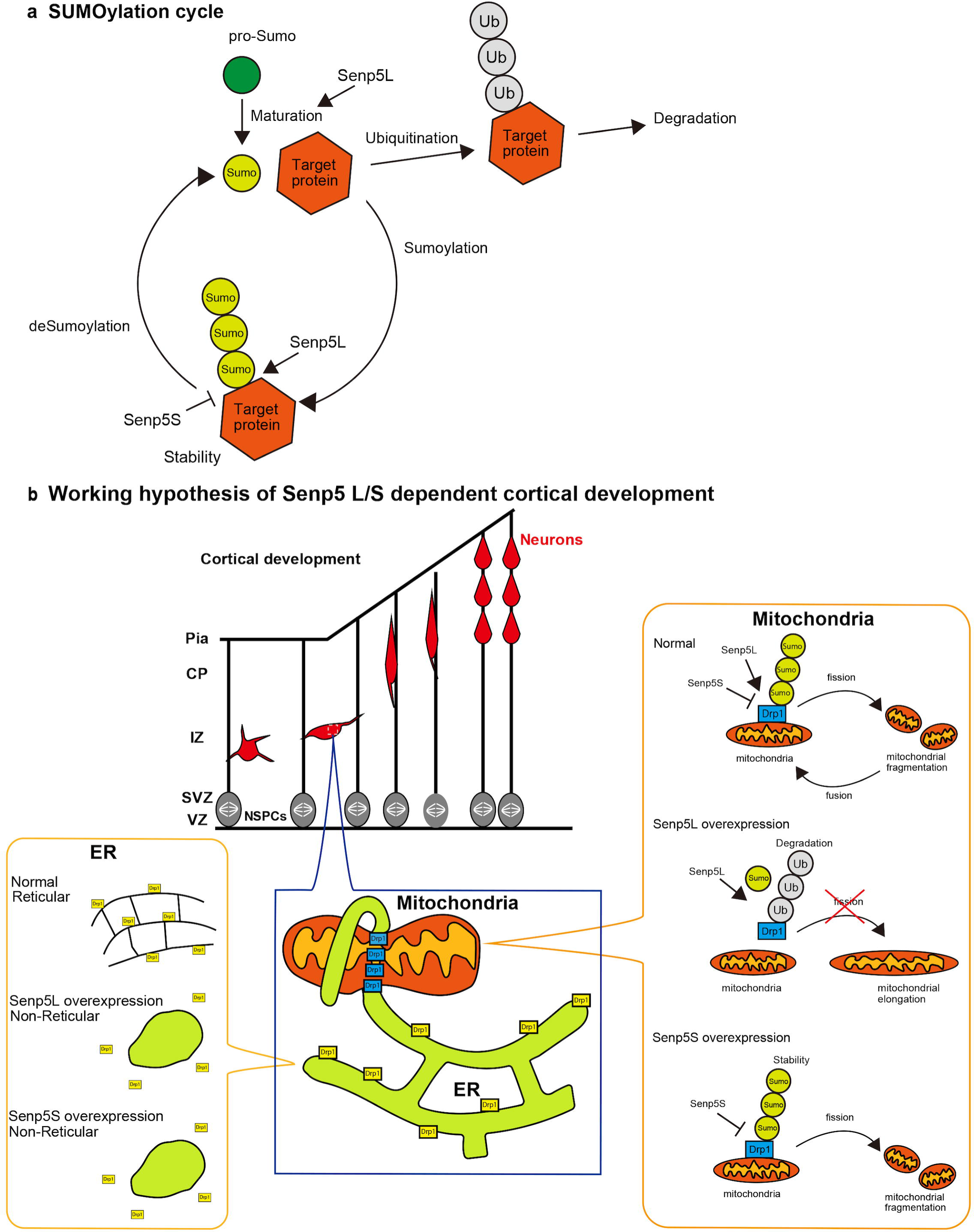
Schematic model of how Senp5L and Senp5S mediate Drp1 SUMOylation. (a) Shown is the SUMOylation cycle and the roles of Senp5L/5S. Senp5L removes SUMO from the target proteins (deSUMOylation) and Senp5S promotes SUMOylation, probably due by substrate competition with other Senps, including Senp5L. Senp5L also is known to cleave pro-SUMO to produce the mature and conjugatable form of SUMO. (b) Hypothetical function of Senp5L/S in regulating cortical development. A tuned balance between Senp5L and Senp5S expression is indispensable for neuronal polarization and cortical organization. Overexpression of Senp5L accelerates deSUMOylation and ubiquitination of Drp1, and thus mitochondrial fusion. Senp5L overexpression also induces the morphological transformation of ER into the non-reticular form, which reduces the availability of interface sites between the peripheral ER and mitochondria, further promoting mitochondria elongation. However, overexpression of Senp5S enhances Drp1-SUMOylation, resulting in the stabilization of Drp1 and mitochondrial fragmentation. Enhanced SUMOylation due to Senp5S upregulation may inhibit localization of Drp1 to the ER, leading to disruption of the interaction between the ER and mitochondria and a subsequent collapse of the reticular ER.

Like Senp5S, several catalytically inactive splice variants are known to regulate post translational modification. FAK-related and PYK2-related non-kinase inhibit phosphorylation mediated by FAK and PYK2, respectively^30^. Another example is protein-tyrosine phosphatase 1B (PTP1B). The variant of PTP1B lacking a catalytic center promotes phosphorylation by competing with PTP1B to maintain tumor cell survival and proliferation^31^. In addition, a catalytically inactive form of TRE/ubiquitin specific protease (UPS6) has been identified, although it remains unclear whether the short isoform and the full-length USP6 competitively regulate the ubiquitination of target proteins^32, 33^. These findings support competitive regulation by catalytically active and inactive splice variants as a common strategy for tuning the post translational modification state of the target molecule. In the present study, we demonstrated the physiologically crucial role of this mechanism, in which Senp5 isoforms had an opposite activity on Drp1 modification and competitively regulated mitochondrial dynamics, thus allowing tight control of neuronal differentiation and corticogenesis.

We showed that Senp5L and Senp5S had opposite effects on SUMO conjugation to Drp1. Senp5L promoted deSUMOylation, while Senp5S facilitated SUMOylation. SUMO3 deconjugation/conjugation from/to Drp1 mediated by each isoform caused mitochondrial elongation and fragmentation, respectively (Fig. 7b). We proposed two possible, non-mutually exclusive, mechanisms for Senp5-dependent control of mitochondrial dynamics. One was through Drp1 ubiquitination, while the other was through Drp1 localization and ER tubulation. Previous studies suggested that Drp1 was stabilized by SUMO1-conjugation, and SUMO1-ylated Drp1 promoted mitochondrial fragmentation^21^. This allowed us to speculate that deSUMOylated Drp1 is a labile form, the formation of which might be quickly followed by protein degradation. Indeed, we showed that Senp5L overexpression enhanced the ubiquitination of Drp1, which led to a lower rate of mitochondrial fission in Senp5L expressing cells. In contrast, Senp5S is likely to stabilize Drp1 by promoting SUMOylation, resulting in mitochondrial fragmentation (Fig. 7b).

Drp1 is localized to the ER and mitochondria^17^. Drp1 distributed around the ER induced ER tubulation independent of its GTPase activity. This structural change facilitated contacts between the ER and mitochondria, leading to mitochondrial fragmentation^17^. Binding between Drp1 and phosphatidic acid (PA) played an essential role in Drp1-mediated ER tubulation, which required four lysines in the D-octadecapeptide sequence^17^. Here, we used Drp1-4KR to demonstrate ER de-structuration, consistently showing the involvement of the four lysines in ER tubulation. Because arginine was used to substitute for lysine rather than alanine, it was unclear whether PA binding was inhibited in our experimental condition. Drp1-4KR would have retained the binding ability if it bound to PA electrostatically. How does SUMOylation regulate the ER tubulation and mitochondrial dynamics? One possible scenario is that Senp5 isoforms control the intracellular distribution of the Drp1 pool as the regulator of SUMOylation level of Drp1. In this scenario, a higher level of Drp1-SUMOylation caused by Senp5S shifts the localization of Drp1 from the ER to mitochondria, resulting in the collapse of ER tubulation and mitochondrial fragmentation. Consistently, the variable domain of Drp1 was necessary for the recruitment of Drp1 to the mitochondria^34, 35^, suggesting the involvement of SUMOylation in the intracellular distribution of Drp1. Drp1 exists in the cytosol as a pool of dimers and tetramers and is recruited by the Drp1 receptors Mff or MiD51/MiD49 located on the mitochondrial membrane to produce mitochondrial constriction. MiD51/MiD49 may recruit the Drp1 dimer, while Mff may selectively bond to the higher-order Drp1 complexes^36^, suggesting the existence of Drp1-assembling machinery. In this context, deSUMO/SUMOylation mediated by Senp5L/5S functioned to affect the conformational change and assembly of Drp1 protein, resulting in the recruitment of subpopulations of Drp1 from the cytosol to the mitochondrial fission site.

Senp5 is involved in various cellular aspects including mitosis, differentiation, and apoptosis in Hela cells or hiPS-cardiomyocytes^26, 37, 38^; however, its physiological significance in the brain is unknown. We demonstrated the involvement of Senp5 isoforms in corticogenesis, to some extent, by regulating neurite outgrowth and/or neuronal polarization. Given that both knockdown and overexpression of Senp5L and Senp5S impaired neuronal development, tightly controlled expression balance between the two isoforms is presumed to be important. Mitochondrial fragmentation impairs neurite outgrowth from cortical neurons in culture, and Drp1 which targets mitochondria seems to play crucial roles^39^. One can speculate that Senp5 isoforms exert their effects through SUMOylation/deSUMOylation-dependent targeting of Drp1 to mitochondria as discussed above. We also showed that Senp5 isoforms disturbed neuronal polarization. Although it is unclear whether the defects in polarity formation are a result of impaired neurite outgrowth, axon specification can be controlled by mitochondrial dynamics^40^ and thus by Senp5 isoforms.

There is another possible explanation for how Senp5 isoforms regulate corticogenesis^18, 19^. Recent *in vivo* imaging studies using mouse embryonic forebrain revealed that NSPCs have an increased ability to undergo mitochondrial fusion, whereas some daughter cells that acquire a high level of mitochondrial fission are destined to be neurons^18^. Therefore, neuronal differentiation can be regulated by mitochondrial dynamics. Considering the expression of Senp5 isoforms in the neuronal lineage from the early to late stage of brain development, Senp5 may be involved in broad aspects of neurogenesis that are closely linked with Drp1 and consequent mitochondrial dynamics.

The expression levels of the Senp5 isoforms in the brain changed as development proceeded; Senp5S expression was drastically upregulated in postnatal and adult brains. Our previous immunoelectron microscopy study revealed that in the adult brain, Senp5 protein localizes to presynaptic terminals, postsynaptic spines, and mitochondria^27^, suggesting a functional relevance of Senp5S to synaptogenesis and synaptic transmission in the mature neuron. Since we confirmed the expression of the SENP5S isoform in humans (UniProt number; Q96HI0-2), which lacks His-646 and Asp-663 in the C-terminal catalytic triad^41^, it is likely that SENP5S plays an evolutionarily conserved role in the development and maintenance of the mammalian nervous system. SUMOylation promotes the stability of several target proteins by antagonizing their ubiquitination^21, 42–44^. An imbalance between SUMOylation and deSUMOylation is implicated in various neurodegenerative diseases in humans such as Alzheimer’s disease (AD), amyotrophic lateral sclerosis (ALS), Parkinson’s disease (PD), and developmental abnormalities^45^. In AD, the enhanced SUMOylation of tau was detected in cerebral cortex. SUMOylation inhibited ubiquitination-mediated tau degradation, leading to the accumulation of insoluble aggregates of tau in the AD brain^46^. Similarly, SUMOylation is involved in SOD1 aggregation in ALS and α-Synuclein aggregation in PD^42, 47^. In addition, several genes associated with PD, such as α-Synuclein, Pink1, and Parkin, control mitochondrial function and morphology. If Senp5L and 5S function in the SUMOylation of these proteins in the adult brain, Senp5 would be a possible therapeutic target for degenerative disorders.

## METHODS

### Animals

Experiments were approved by the Committee on the Ethics of Animal Experiments of the Waseda University. ICR mice (SLC Japan) were maintained on a 12-hour light–dark cycle and had continuous access to food and water. The date of conception was established by the presence of a vaginal plug and was recorded as embryonic day zero (E0). The day of birth was designated as P0.

### Cell culture and transfection

HEK293T and Neuro2a cells were cultured in Dulbecco’s modified eagle medium (DMEM) (Fujifilm-Wako) containing 10% fetal bovine serum (FBS) (Biowest), penicillin/streptomycin, and 2 mM L-glutamine (Thermo) in a humidified atmosphere of 5% CO_2_/95% air at 37°C. Primary NSPCs were isolated from the E12 telencephalon and cultured as previously described^48^. Neuron cultures were prepared from the cerebral cortex of E16-17 mice. Cortices were dissociated using 0.25% trypsin followed by trituration. Cells were plated on coverslips coated with 0.01% poly-D-lysine (PDL) and incubated in Neurobasal medium (Thermo) supplemented with the B-27 (Thermo) and GlutaMAX (Thermo) in a humidified atmosphere of 5% CO_2_/95% air at 37°C. Neurons were processed for immunocytochemistry or western blotting after 3 days *in vitro*. Cultured cell lines were transfected with plasmids using PEI MAX (Polysciences) complexes with a DNA to PEI MAX ratio of 1:3 w/w as previously described^49^. Mouse NSPCs and primary cortical neurons were electroporated using the NEPA21 Electroporator (Nepagene) according to the manufacturer’s instructions.

### Cloning of mouse Senp5 cDNAs

Total RNA was isolated from C57BL/6 mice brains at 10 weeks old using the RNAiso Plus reagents (Takara Bio). First strand cDNA was synthesized using the PrimeScript II cDNA Synthesis Kit (Takara Bio) following the manufacturer’s protocol. The cDNAs encoding the Senp5 open reading frame (ORF) were amplified by Q5 High-Fidelity DNA Polymerase (NEB Japan) with primer sets based on the predicted sequences of the mouse Senp5 isoforms (GenBank accession numbers: Senp5 short isoform 1 [Senp5S], XR_003951796; Senp5 short isoform 2 [Senp5S2], NM_001357088 and XR_004939194). The primers were as follows: forward primer (common to both isoforms), 5′-TCTCGAGCCAGTTCTCATTATGCATCAG-3′; reverse primer for short form 1, 5′-ATGGTACCTTCAGTACAAAGGGAGACAG-3′. PCR products were sequenced and cloned into the XhoI/KpnI sites of the pEGFP-C2 expression vector (Takata Bio). Isolated cDNA clones were designated as Senp5S and Senp5S2. The ORF nucleotide sequences were deposited with the GSDB, DDBJ, EMBL, and NCBI under accession numbers LC619058 (Senp5S) and LC619059 (Senp5S2).

### Quantitative PCR

Total RNAs from the mouse forebrain were reverse transcribed using the PrimeScript II first strand cDNA Synthesis Kit (Takara Bio) with the random hexamer primer. PCR was performed using the TB Green Ex Taq II Mix (Takara Bio) and the Thermal Cycler Dice Real Time System (Takara Bio) according to the manufacturer’s protocol. The primers were as follows: Senp5L (NM_001357087.1); forward primer, 5′-AGACAGCTGGTAACCAAAGGCT-3′; reverse primer, 5′-TCCGGCTGGAGAGTGTCACAGT-3′, Senp5S; forward primer, 5′-AGACAGCTGGTAACCAAAGGCT-3; reverse primer, 5-GTACAAAGGGAGACAGAATACCTTC-3′, β-actin; forward primer, 5′-GGCTGTATTCCCCTCCATCG-3′; and reverse primer, 5′-CCAGTTGGTAACAATGCCATGT-3′. PCR products were loaded on a 1% agarose gel and visualized to confirm the specificity of each primer set.

### Plasmids

pEGFP-Senp3 and pEGFP-Senp5L were provided by Dr. T. Nishida (Mie University, Mie, Japan)^50, 51^. pIRES2-mCherry-Senp5L and pIRES2-mCherry-Senp5S, from which the Senp5 and mCherry genes were translated from a single bicistronic mRNA, were generated through insertion of the Senp5 ORF into pIRES2-mCherry. The pIRES2-mCherry vector was synthesized by replacing the AcGFP part of pIRES2-AcGFP1 (Takara Bio) with the mCherry sequence. pEGFP and pmCherry expression vectors were purchased from Takara Bio. pMT-mKO1, which expressed mitochondria-targeted CoralHue monomeric orange fluorescent protein (Kusabira Orange; MBL/Amalgaam brand, AM-V0221M), was used for mitochondrial imaging. mCherry-Sec61B (Addgene plasmid # 121160, deposited by Dr. Mayr)^52^ was used for ER imaging. Myc-Drp1, cloned from mouse, was provided by Dr. M. Kengaku (Kyoto University, Japan)^53^. YFP-Drp1 (human isoform 3) and the YFP-Drp1-4KR mutant were provided by Dr. Chun Guo (University of Sheffield, UK)^25^. Wild-type Drp1 or non-SUMOylatable Drp1-4KR were subcloned into a pCAGGS vector to construct the pCAG-YFP-Drp1 and pCAG-YFP-Drp1-4KR plasmids. The CAG promoter was used to facilitate upregulation of Drp1 and Drp1-4KR *in vivo*. Plasmid constructs containing short hairpin RNA (shRNA) cassettes in the pRFP-C-RS vector were purchased from OriGene Technologies. Plasmid-transfected cells expressed shRNA, which was controlled by the U6 polymerase III promoter and RFP fluorescent protein^27^. The silencing effect of the Senp5 shRNAs was established in our previous report. The shRNA sequences were as follows: 5′-GCACTACCAGAGCTAACTCAGATAGTACT-3′ (scrambled non-targeting control), 5′-GCAGGTGAAGCATTTCCAGTGTAATAAGG-3′ (Senp5 shRNA#1), and 5′-CTGGAGCAGACAAATCTGTGAGTAGTGTA-3′ (Senp5 shRNA#2). The shRNA #1 and shRNA #2 target both isoforms of Senp5 mRNA^27^.

### In utero electroporation

*In utero* electroporation was performed as described previously^54^. Pregnant mice were anesthetized with medetomidine, midazolam, and butorphanol. Plasmid DNA solution was injected into the lateral ventricle of E14.5 mice through the uterus wall and electroporated using the NEPA21 electric-pulse generator (Nepagene) according to the manufacturer’s instructions. Embryos were dissected out from the uteri two days after electroporation. They were then perfused transcardially with saline solution and 4% paraformaldehyde (PFA) in 0.1 M phosphate buffer (pH, 7.4).

### Immunoprecipitation and immunoblotting

For immunoprecipitation (IP), cells were washed twice in ice-cold phosphate-buffered saline (PBS) and lysed in ice-cold lysis buffer containing 50 mM Tris-HCl (pH, 7.4), 150 mM NaCl, 2 mM EDTA, 1% NP-40, 20 mM N-ethyl-maleimide (NEM), and protease inhibitors (Complete Mini Protease Inhibitor Cocktail, Sigma-Aldrich) for 30 min at 4°C.

For the ubiquitination assay, cells pretreated with or without 5 mM MG132 for 3 h were washed twice with PBS and lysed in denaturing RIPA buffer containing 1% sodium dodecyl sulfate (SDS), 5 mM EDTA, 10 mM dithiothreitol (DTT), 20 mM NEM, and protease inhibitors. Cell lysates were heated for five min at 95°C, followed by a 10-fold dilution with lysis buffer. Lysates were centrifuged at 15,000 rpm for 10 min. The supernatants were incubated with primary antibody for 60 min. The supernatants with primary antibody were coupled with protein A/G plus agarose IP beads (Santa Cruz Biotechnology) overnight at 4°C. After centrifugation, the beads were washed four times with 0.1% NP40 in PBS. The bound proteins were dissolved with 2× sample buffer (125 mM Tris-HCl, pH 6.8, 4% SDS, 10% sucrose, and 0.01% bromophenol blue). IP samples were resolved on 8% or 10% SDS-PAGE gels and transferred onto Immobilon-P membranes (Merck Millipore) using a semidry transfer apparatus. The membranes were blocked using Blocking One (Nacalai Tesque) for 25 min or 5% skim milk for 60 min at room temperature. After washing with 0.1% Tween 20 in Tris-buffered saline (TBST), the membranes were incubated with primary antibody against GFP (1:2,000, GeneTex, GTX113617), HA (1:10,000, MBL, TANA2), Myc (1:7500, MBL, My3), Senp5 (Proteintech, 19529-1-AP), or α-tubulin (1:2,000, MBL, #PM054; 1:5000, Proteintech, 11224-1-AP) diluted in TBST, for 60 min at room temperature. Membranes were subsequently incubated with horseradish peroxidase (HRP)-conjugated anti-rabbit IgG (1:20,000; Cytiva, NA934) or anti-mouse IgG (1:20,000; Cytiva, NA931) for 60 min at room temperature. Blots were visualized using the Immobilon western HRP substrate (Merck Millipore) and the Fusion Solo S (Vilber Lourmat) or LAS-3000 luminescent image analyzer (FujiFilm). ImageJ was used to quantify band densities.

### Immunocytochemistry

Immunocytochemistry was performed as described previously^48^. Cortical neurons and cultured cells were fixed with 4% PFA, permeabilized with 0.1% Triton X-100, and blocked with 10% normal goat serum at room temperature for 60 min. Cells were then incubated with anti-tagRFP (1:1,000, Evrogen, AB233), anti-SMI312 (1:1,000, Biolegend, 837904), and anti-HA (1:1000, MBL, TANA2) at room temperature for 60 min. Primary antibody binding was visualized using the Alexa Fluor-conjugated secondary antibody (1:1500, Thermo). Immunofluorescence images were acquired using 40× objective lenses (NA, 0.75; Zeiss) and a CCD camera (AxioCam MRm, Zeiss) mounted on an inverted microscope (Axio Observer, Zeiss).

### Immunohistochemistry

Embryos were fixed by immersion or perfusion in the cardiac ventricle with 4% PFA in 0.1 M phosphate buffer (pH 7.4), followed by post-fixation overnight at 4°C. Fixed embryo brains were cryoprotected in PBS with 30% sucrose overnight at 4°C and embedded in an optimal cutting temperature compound (Sakura Finetek). Frozen sections were cut at a thickness of 14 μm using a cryostat and were collected on MAS coated glass slides (Matsunami Glass). For antigen retrieval, tissue sections were heated at 90 to 95°C for 10 min in 10 mM sodium citrate buffer (pH, 6.0) using a microwave oven, followed by treatment with 0.3% H_2_O_2_ in PBST (0.1% Triton X-100 in PBS) for 40 min at 25°C to quench endogenous peroxidase activity. Sections were blocked for an hour in 5% normal horse serum in PBST and incubated overnight at 4°C with primary antibodies diluted in the same blocking solution. Sections were incubated with HRP-polymer linked anti-rabbit IgG reagent (ImmPRESS; Vector Lab) for 2 h. HRP signal was visualized with 0.25 mg/mL diaminobenzidine and 0.03% H_2_O_2_. Each step was followed by four washes with PBST. Free-floating sections were mounted on glass slides, dehydrated, and coverslipped with Entellan New (Merck Millipore). For control sections, primary antibodies were omitted or replaced with normal rabbit serum.

For double immunostaining, frozen sections, treated with antigen retrieval solution, were blocked for 2 h at 4°C with 5% normal goat serum in PBST and incubated with anti-EGFP (1:2000, Aves Labs, GFP-1010), anti-HA (1:500, MBL, TANA2), and anti-Tom20 (1:300, Cell Signaling Technology, D8T4N) antibody overnight at 4°C. After washing with PBST, sections were incubated with Alexa Fluor 488-, Alexa Fluor 555- (Thermo), or DyLight 549- (Jackson ImmunoResearch) conjugated secondary antibodies [F(ab’)_2_ fragment]. After counterstaining with 0.7 nM Hoechst 33342 (Thermo), sections were mounted and imaged using a confocal (FV3000, Olympus) or fluorescence inverted microscope (Axio Observer, Zeiss).

### Analysis of mitochondrial and ER morphology

In mitochondrial morphological analysis, 5–10 z-stack images (1-μm steps) of pMT-mKO1 plasmid-transfected cells were acquired using the confocal microscopy system (FV3000, Olympus). Deconvolution was applied to maximum projection images to clarify the mitochondrial contours. Mitochondrial length was defined as the longitudinal axial length of each mitochondrial fragment, which were traced with cellSens life science imaging software (Olympus). Mitochondrial length was calculated as the mean length of mitochondria within a single cell.

In ER morphological analysis, mCherry-Sec61B-expressing cells were kept in a humidified atmosphere at 37°C using a Stage Top incubator (Tokai-hit). Live cell images were acquired using the confocal microscopy system (FV3000, Olympus). The standard deviation (SD) and mean of the fluorescence intensity were obtained using line-scan analysis by the FV31S-SW (Olympus) in a randomly selected cell region. The ER morphology was defined as reticular when the coefficient of variation (CV) scores (SD/mean) were >0.2.

### Statistics

Statistical analyses were performed using the R package Version 3.4.2. Welch’s *t*-test or the Chi-square test was used to compare two groups. The P value was corrected using the Holm-Bonferroni correction when there were three or more groups. P < 0.05 was considered statistically significant.

## Acknowledgements

The authors would like to thank Dr. C. Guo (The University of Sheffield, UK) for sharing the YFP-Drp1-WT and YFP-Drp1-4KR plasmid, Dr. M. Kengaku for the myc-Drp1, and Dr. T. Nishida (Mie University, Japan) for the EGFP-Senp3 and EGFP-Senp5L. We would like to thank Editage (www.editage.com) for English language editing. We would also thank Mrs. Y. Ishiguro and D. Yasui (Waseda University) for technical assistance.

This work was funded by the Japan Society for the Promotion of Science grants-in-aid [KAKENHI] grant numbers 18K06491 (to H.A.) and 26430 042 (to S.S.) and by Waseda University Grants for Special Research Projects 2018K-352 (to H.A.), 2015K-249, and 2017K-301 (to S.S.).

## Author contributions

All authors have full access to the data of this study and take responsibility for its integrity and accuracy. Study concept and design: S.Y., H.A., and S.S. Acquisition of data: S.Y., A.S., H.A, and S.S. Analysis and interpretation of data: S.Y., A.S., H.A, and S.S. Drafting of the manuscript: S.Y., H.A, and S.S. Obtained funding: H.A and S.S.

