## Supplemental text for "Drp1 SUMO/deSUMOylation by Senp5 isoforms influences ER tubulation and mitochondrial dynamics to regulate brain development"

**SUPPLEMENTAL INFORMATION**

**Supplementary Figure 1. SUMO conjugation to Drp1 adjusts mitochondrial dynamics**

(a, b) HEK293T cells were co-transfected with EGFP-Senp3 or HA-SUMO1 together with pMT-mKO1, followed by immunocytochemistry with an anti-HA (*green*). Higher magnification of the boxed areas shows pMT-mKO1^+^ mitochondrial morphology (*white*). Arrows (*yellow*) denote fragmented mitochondria. (b) The box and whisker plots summarize the mitochondrial length (μm). Numbers in parentheses indicate the numbers of cells measured. ***, P < 0.001; Welch’s t-tests with Holm-Bonferroni correction. Scale bars, 5 μm.

**Supplementary Figure 2. Drp1 deSUMOylation promotes NEM dependent ubiquitinations**

HEK293 cells expressing EGFP, EGFP-Senp5L, or EGFP-Senp5S together with Myc-Drp1 and HA-SUMO3 were subjected to immunoprecipitation with an anti-HA antibody with ± N-ethyl-maleimide (NEM), followed by immunoblotting with anti-HA or anti-Myc antibodies.

**Supplementary Figure 3. Senp3 expression in the developing cortex**

(a–c) Coronal sections of the cerebral cortex at E13.5 (a) and E16.5 (b) immunostained with an anti-Senp3 antibody. (c) Higher magnification of the area of the cerebral cortex surrounding the lateral ventricle (lv) in b. Scale bars, 100 μm in (a, b); 25 μm in (c). lv, lateral ventricle; CP, cortical plate; IZ, intermediate zone; VZ, ventricular zone.

**Supplementary Figure 4. Validation of *Senp5* shRNAs against Senp5S**

Neuro2a cells were transfected with pCAG-EGFP-Senp5S and two different mouse *Senp5* shRNA constructs (shRNA #01 and shRNA #02) or a non-targeting control shRNA, followed by immunoblotting with an anti-GFP or anti-α-tubulin antibody.
