## Supplementary figures and images for "Drp1 SUMO/deSUMOylation by Senp5 isoforms influences ER tubulation and mitochondrial dynamics to regulate brain development"

### Supplemental Figure 1

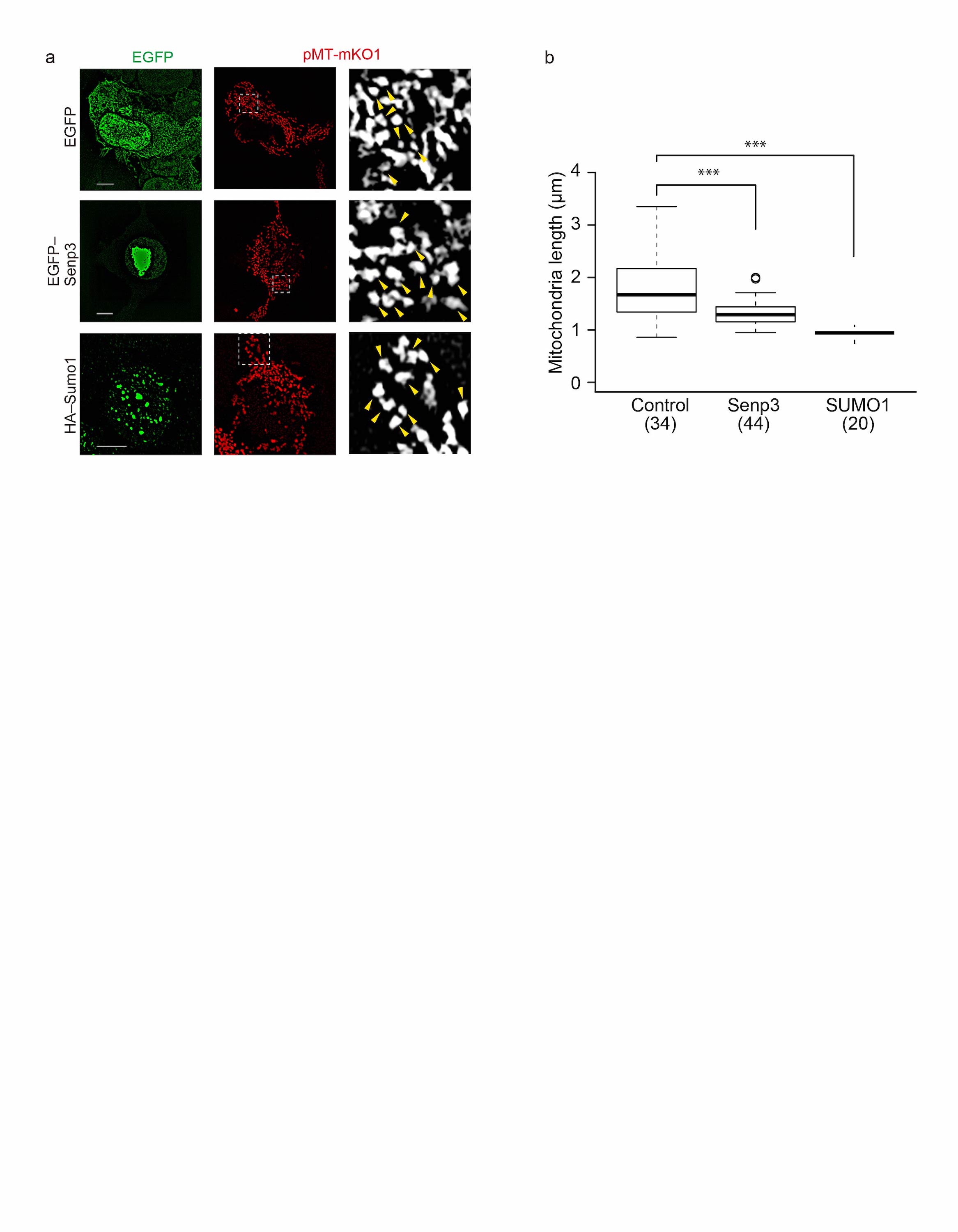

### Supplemental Figure 2

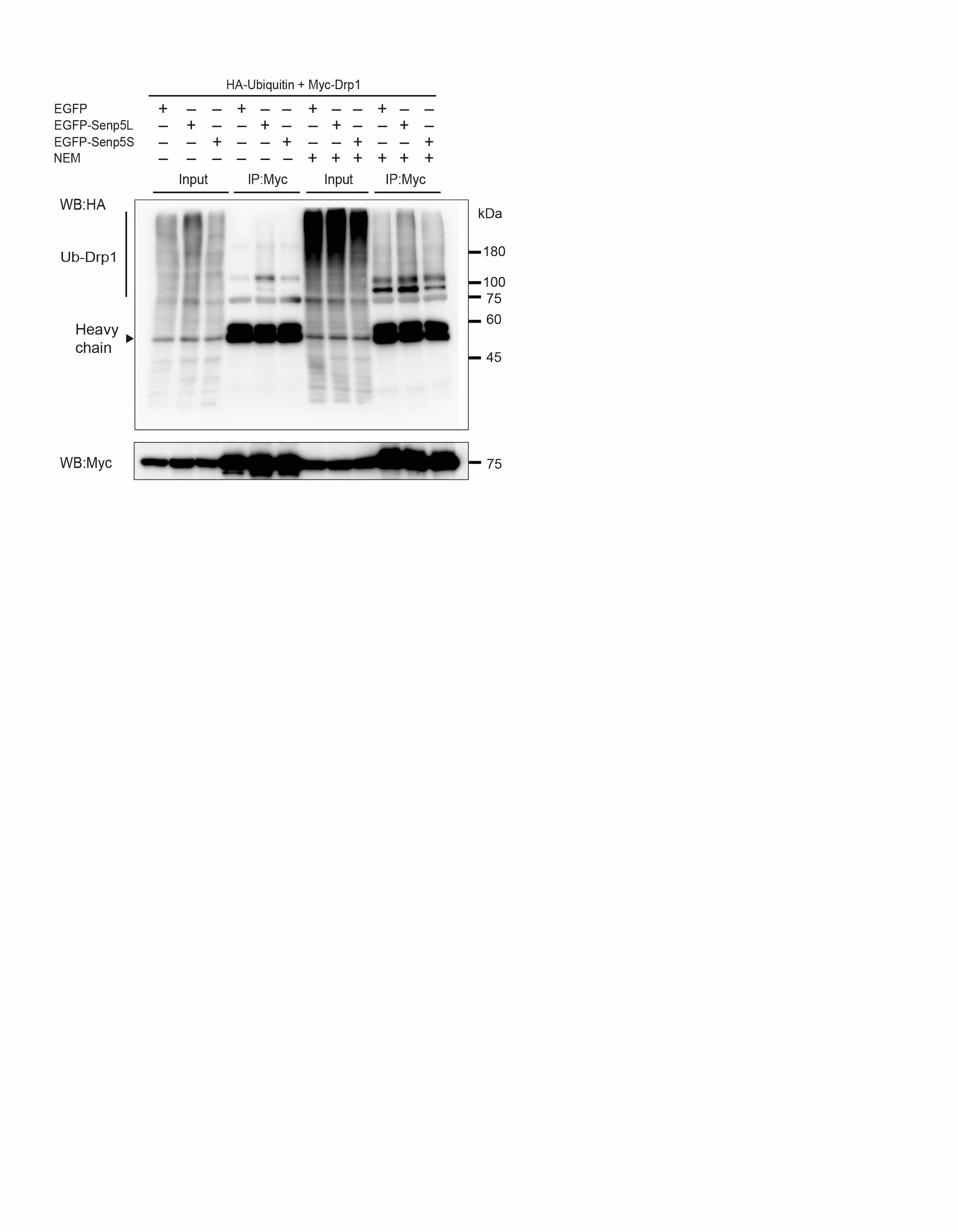

### Supplemental Figure 3

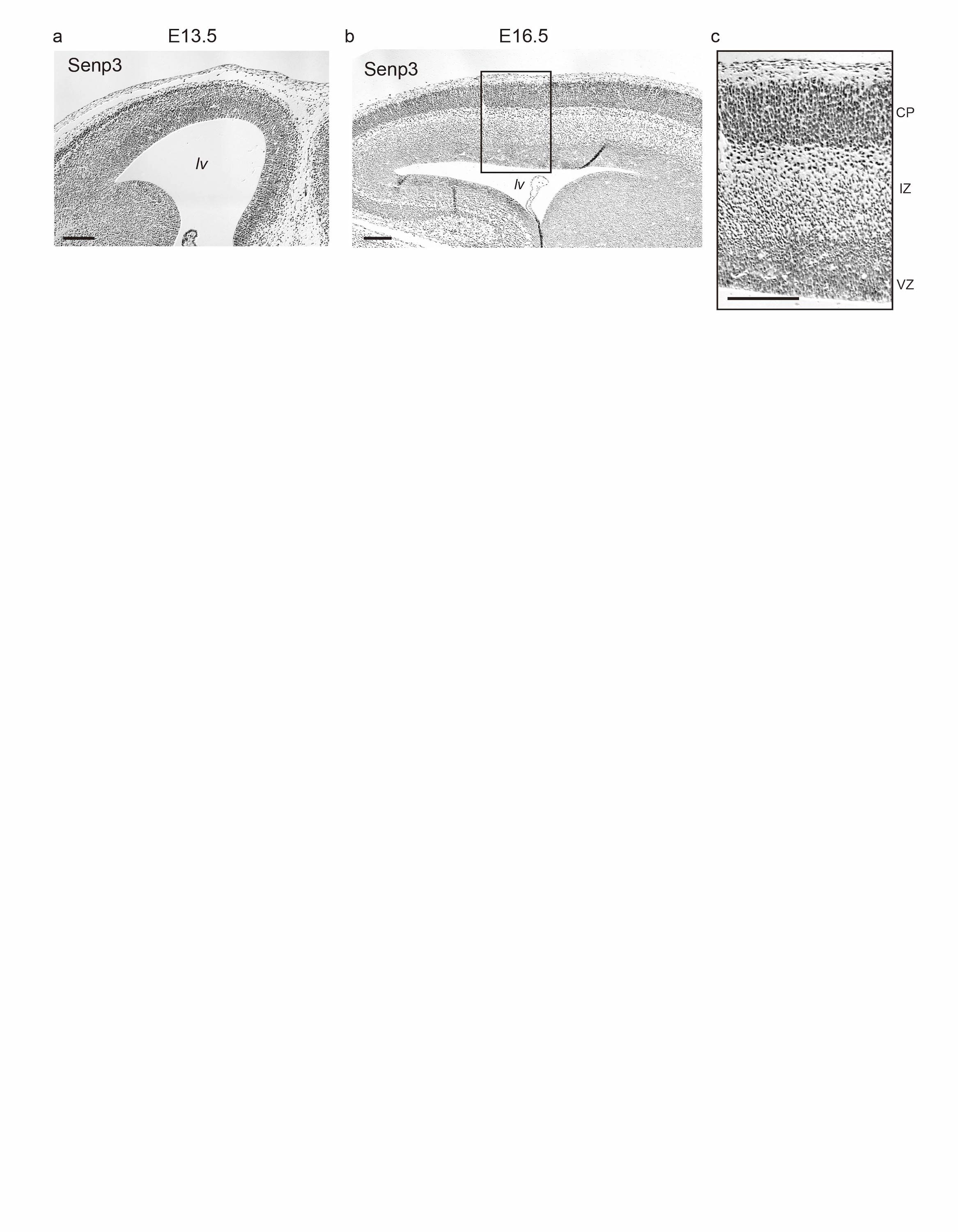

### Supplemental Figure 4

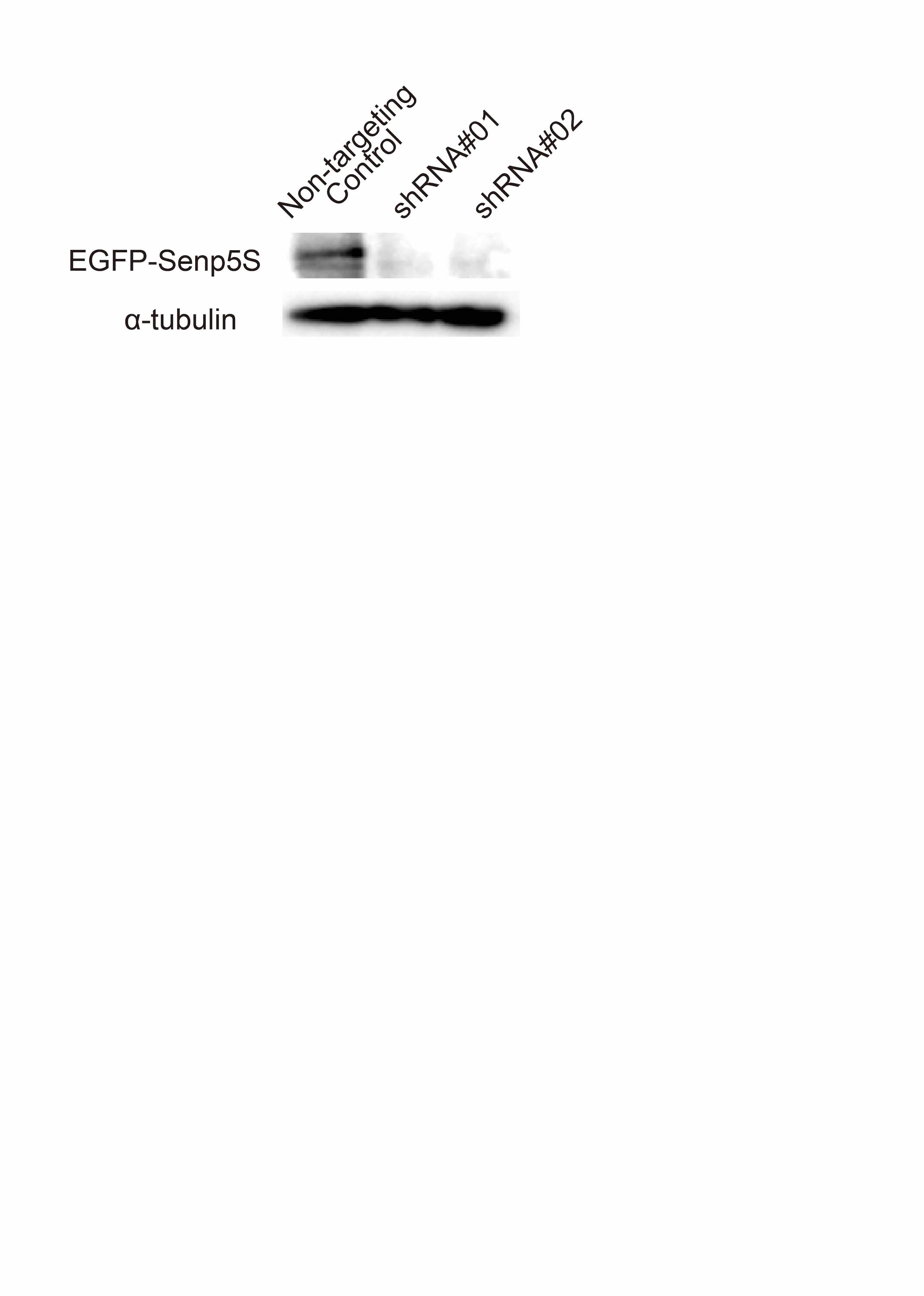
